# *Pseudomonas aeruginosa* performs chemotaxis to serotonin, dopamine, epinephrine, and norepinephrine

**DOI:** 10.1101/2024.12.05.626837

**Authors:** Elizabet Monteagudo-Cascales, Andrea Lozano-Montoya, Tino Krell

## Abstract

Bacteria use chemotaxis to move to favorable ecological niches. For many pathogenic bacteria, chemotaxis is required for full virulence, particularly for the initiation of host colonization. There do not appear to be limits to the type of compounds that attract bacteria, and we are just beginning to understand how chemotaxis adapts them to their lifestyles. Quantitative capillary assays for chemotaxis show that *P. aeruginosa* is strongly attracted to serotonin, dopamine, epinephrine, and norepinephrine. Chemotaxis to these compounds is greatly decreased in a mutant lacking the TlpQ chemoreceptor, and complementation of this mutant with a plasmid harboring the *tlpQ* gene restores wild-type-like chemotaxis. Microcalorimetric titrations of the TlpQ sensor domain with these four compounds indicate that they bind directly to TlpQ. All four compounds are hormones and neurotransmitters that control a variety of processes and are also important signal molecules involved in the virulence of *P. aeruginosa*. They modulate motility, biofilm formation, the production of virulence factors and serve as siderophores that chelate iron. Therefore, chemotaxis to these four compounds is likely to alter *P. aeruginosa* virulence. Additionally, we believe that this is the first report of bacterial chemotaxis to serotonin and dopamine. This study provides an incentive for research to define the contribution of chemotaxis to these host signaling molecules to the virulence of *P. aeruginosa*.

## Importance

Serotonin, dopamine, epinephrine, and norepinephrine have been shown to be central signal molecules that contribute to the virulence of *P. aeruginosa* in a number of animal model systems. Chemotaxis to these compounds is thus likely to modulate virulence. This report provides insight into the forces that have driven the evolution of the bacterial chemotaxis system and will motivate studies to establish whether other pathogens are also attracted to these compounds. The observation of chemotaxis to serotonin and dopamine expands the range of bacterial chemoattractants. Interference with chemoreceptor function is thus suggested to be an alternative strategy to fight bacterial pathogens and may promote the development of novel antimicrobial agents.

Bacteria use chemotaxis to move to sites that are favorable for growth and survival. A chemotactic response is typically initiated by the binding of a chemoeffector to a chemoreceptor, stimulating chemosensory pathways and ultimately leading to chemotaxis. There do not appear to be any limits to the chemical properties or structure of chemoeffectors. The chemosensory capacities of a bacterium are a reflection of its lifestyle. As the chemoeffectors recognized by most chemoreceptors are unknown, we are only beginning to understand the link between chemosensory repertoire, physiology, and lifestyle. For pathogens with very different lifestyles, chemotaxis is often essential for full virulence, and particularly for the initiation of host colonization (1). *Pseudomonas aeruginosa* is among the most serious human pathogens. It kills about half a million people annually (2). The genome of the *P. aeruginosa* strain PAO1 encodes 26 predicted chemoreceptors, and the corresponding chemoeffectors have been identified for about half of them (3). Chemotaxis is required for full virulence of *P. aeruginosa* (4–6). The catecholamines dopamine, epinephrine, serotonin, and norepinephrine are signal molecules that act as hormones and neurotransmitters to control a variety of cellular processes in animals (7). They also exert signaling functions in plants (8) and bacteria (9), and they play major roles in inter-kingdom signaling (10). To determine whether *P. aeruginosa* performs chemotaxis to these compounds, we conducted quantitative capillary chemotaxis assays. Strong chemoattractant responses were observed for all compounds (Fig. 1A).

**Fig. 1).**
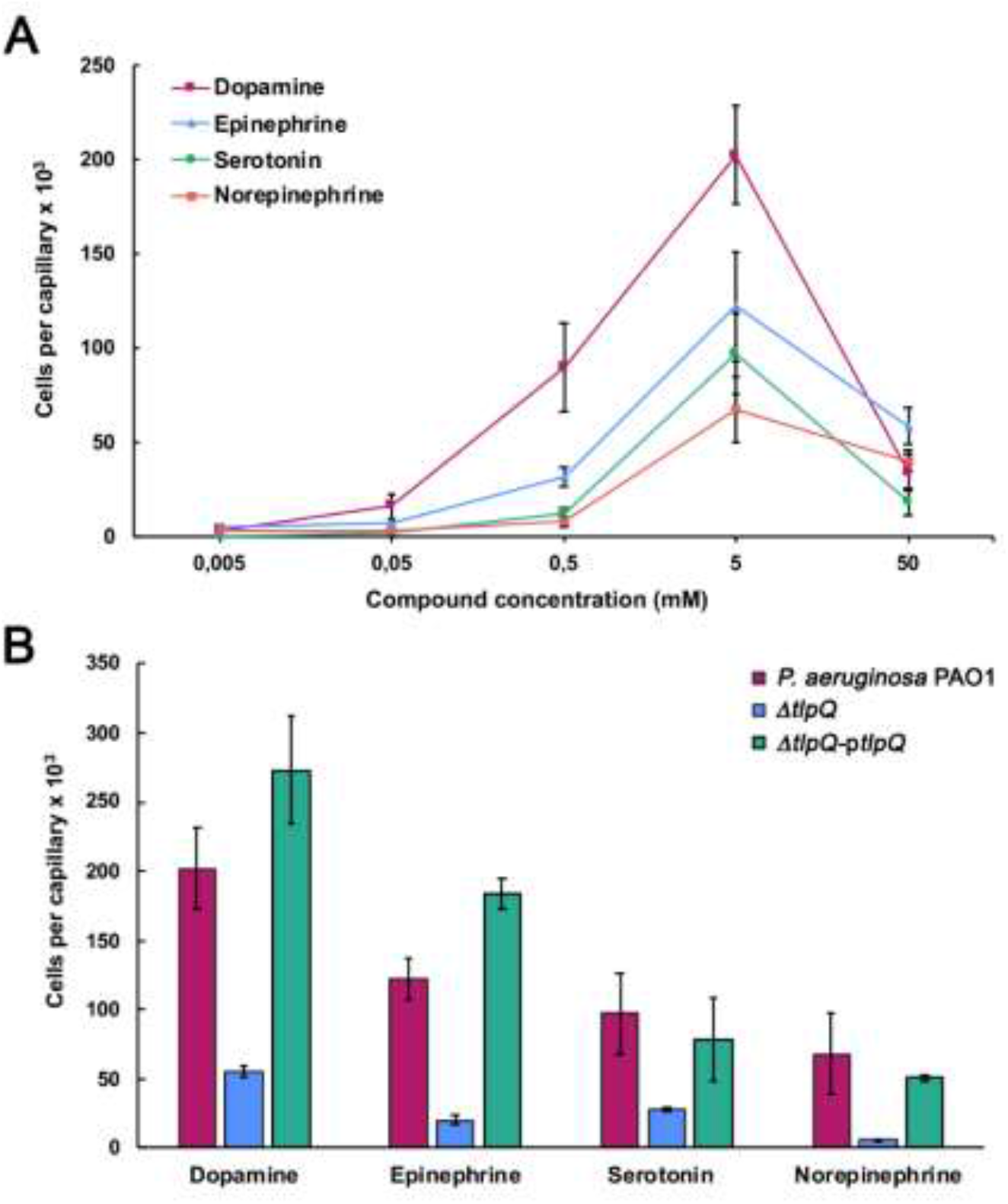
Chemotactic responses to dopamine, epinephrine, serotonin, and norepinephrine of *Pseudomonas aeruginosa* PAO1 and a mutant deficient in the TlpQ chemoreceptor. **A)** Quantitative capillary chemotaxis assays of the wild-type strain to different chemoeffector concentrations. Data were corrected for the number of bacteria that swam into buffer-containing capillaries (3107 ± 821). **B)** Responses to 5 mM of these four neurotransmitters by the wild-type strain, the Δ*tlpQ* mutant, and the Δ*tlpQ* mutant complemented with the p*tlpQ* plasmid. Data were corrected for the number of bacteria that swam into buffer-containing capillaries (4071 ± 1536 for wt; 4143 ± 604 for Δ*tlpQ*, and 2071 ± 371 for Δ*tlpQ*-p*tlpQ*). Data are the means and standard deviations from at least three biological replicates conducted in triplicate.

The threshold for chemotaxis was at capillary concentrations of 50 μM for dopamine and epinephrine and 500 μM for serotonin and norepinephrine (Fig. 1A). The maximum responses were seen at a capillary concentration of 5 mM and ranged from 65,000 to 200,000 cells per capillary. These responses are similar to those seen with other chemoattractants like nitrate (11), and superior to those seen with malate (12) or α-ketoglutarate (13). Whereas bacterial chemotaxis to epinephrine and norepinephrine has been observed previously (14, 15), we believe this is the first report of chemotaxis to dopamine and serotonin.

To identify the chemoreceptor(s) that detect these four molecules, we conducted chemotaxis assays using a quadruple mutant in the *pctA, pctB, pctC*, and *tlpQ* chemoreceptor genes. As shown in Fig. S1, chemotactic responses to all four compounds were lower than in the wild-type strain. Since PctA, PctB, and PctC are known to be chemoreceptors for amino acids (16), we performed chemotaxis assays using a *tlpQ* mutant. Responses to all four molecules were significantly decreased, suggesting that TlpQ is their primary chemoreceptor (Fig. 1B). Complementation of the mutant with a plasmid harboring the *tlpQ* gene resulted in wild-type-like chemotaxis to all four compounds (Fig. 1B). Control experiments showed that the responses of the *tlpQ* mutant and the wild-type strain to casamino acids are comparable (Fig. S2).

Chemoreceptors can be activated by the binding of chemoeffectors or chemoeffector– loaded solute binding proteins to the chemoreceptor ligand binding domain (LBD). To determine the mode of TlpQ activation, we purified the TlpQ-LBD and subjected it to microcalorimetric titrations with the different compounds. Whereas the titration of buffer with dopamine resulted in small and uniform peaks, representing dilution heats, titration of TlpQ-LBD resulted in exothermic heat changes that diminished as protein saturation advanced (Fig. 2A).

**Fig. 2).**
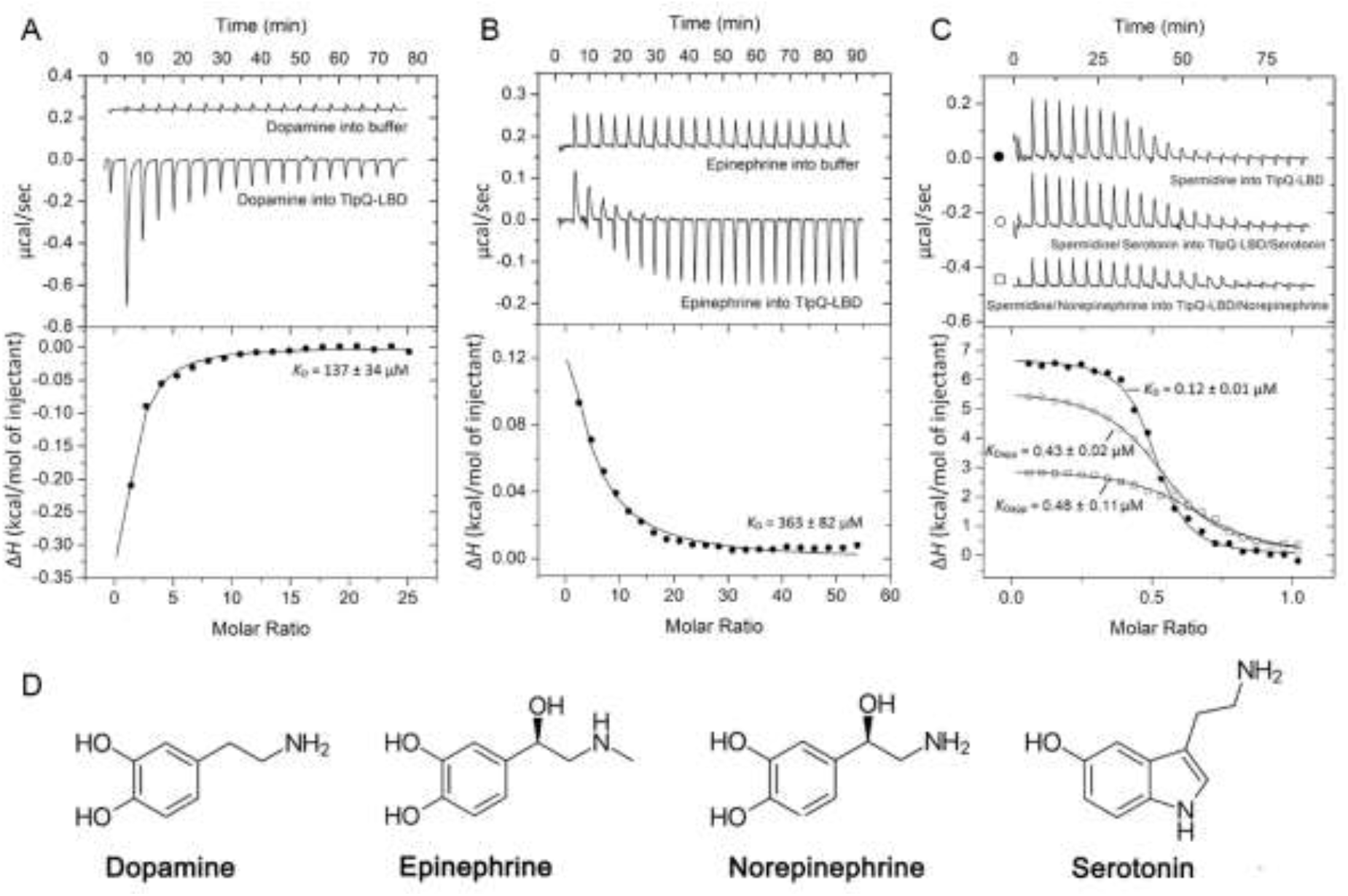
Microcalorimetric titrations of dopamine, epinephrine, serotonin, and norepinephrine to the TlpQ-LBD. **A)** Titration of 80 μM TlpQ-LBD with aliquots of 10 mM dopamine. **B)** Titration of 40 μM TlpQ-LBD with aliquots of 10 mM epinephrine. **C)** Competition assays: Titration of 18 μM TlpQ-LBD with aliquots of spermidine in the absence and presence of 10 mM serotonin or norepinephrine. Upper panels: Raw titration data. Lower panels: Integrated, concentration-normalized and dilution-heat-corrected raw data fitted with the “one-binding-site” model of the MicroCal version of ORIGIN. **D)** Structure of the TlpQ ligands identified.

The dissociation constant for dopamine was 137 μM. Titration with epinephrine also showed binding (*K*_D_=363 μM, Fig. 2B). Initial titrations with serotonin and norepinephrine gave indications of binding, but with an affinity lower than that for dopamine and epinephrine. The interaction of serotonin and norepinephrine with the TlpQ-LBD could be better visualized using competition assays. Previous studies have shown that TlpQ also binds polyamines like spermidine (17). The titration of TlpQ-LBD with spermidine resulted in a *K*_D_ of 120 nM (Fig. 2C). When the TlpQ-LBD was titrated with spermidine in the presence of 10 mM serotonin or norepinephrine, a significant reduction in binding was observed (Fig. 2C), with lower changes in binding enthalpy (Δ*H*_app_) and increased dissociation constants (*K*_Dapp_) in the presence of either serotonin or norepinephrine compared to titration without competitor. Thus, serotonin and norepinephrine compete with spermidine for binding to the TlpQ-LBD. We conclude that TlpQ is activated by direct binding of dopamine, epinephrine, serotonin, and norepinephrine.

Multiple studies show that these four catecholamines are central signal molecules in *P. aeruginosa*. Dopamine (18) and serotonin (19) act as exogenous quorum-sensing molecules that regulate the production of a wide range of virulence factors, motility, and biofilm formation. Epinephrine and norepinephrine increase pyoverdine and pyocyanine production (20) as well as adhesion and biofilm formation (21). Furthermore, these four catecholamines serve *P. aeruginosa* as siderophores (22, 23). Experimentation using different animal models show that the four catecholamines modulate *P. aeruginosa* virulence (19, 21, 24, 25).

Taken together, the directed movement of *P. aeruginosa* in gradients of these compounds is likely to contribute to its virulence. This study thus forms the basis for exploring the effect of chemotaxis to catecholamines on *P. aeruginosa* virulence. It also suggests that other motile bacterial pathogens should be tested for chemotaxis to these compounds.

## Materials and Methods

Provided in the Supplementary Material.

## Acknowledgements

We are indebted to Dr. Michael Manson for critical reading and editing the manuscript. This study was supported by grants from the Spanish Ministry for Science and Innovation/*Agencia Estatal de Investigación* 10.13039/501100011033 (grants PID2020-112612GB-I00 and PID2023-146216NB-I00 to TK) and the Consejo Superior de Investigaciones Científicas (grant 2024AEP062 to TK).

## Author contribution

EMC: conceptualization, conducted research, analyzed data, wrote revised version of the manuscript; ALM: conducted research, analyzed data, TK: conceptualization, fundraising, wrote manuscript

## References

1. Matilla MA, Krell T. 2018. The effect of bacterial chemotaxis on host infection and pathogenicity. FEMS Microbiol Rev 42:fux052.

2. GBD 2019 Antimicrobial Resistance Collaborators. 2022. Global mortality associated with 33 bacterial pathogens in 2019: a systematic analysis for the Global Burden of Disease Study 2019. Lancet S0140-6736(22)02185–7.

3. Krell T, Matilla MA. 2024. Pseudomonas aeruginosa. Trends Microbiol 32:216–218.

4. Sheng S, Xin L, Yam JKH, Salido MM, Khong NZJ, Liu Q, Chea RA, Li HY, Yang L, Liang ZX, Xu L. 2019. The MapZ-Mediated Methylation of Chemoreceptors Contributes to Pathogenicity of Pseudomonas aeruginosa. Front Microbiol 10:67.

5. Schwarzer C, Fischer H, Machen TE. 2016. Chemotaxis and Binding of Pseudomonas aeruginosa to Scratch-Wounded Human Cystic Fibrosis Airway Epithelial Cells. PloS one 11:e0150109.

6. McLaughlin HP, Caly DL, McCarthy Y, Ryan RP, Dow JM. 2012. An orphan chemotaxis sensor regulates virulence and antibiotic tolerance in the human pathogen Pseudomonas aeruginosa. PloS one 7:e42205.

7. Savaliya, M., Goerge, J.J. The monoaminergic system in humans. Recent Trends in Science and Technology-2020 (ISBN: 9788192952154) Rajkot, Gujarat, India: Christ Publications, 2020 pp 190–203.

8. Kulma, A., Szopa, J. Catecholamines are active compounds in plants. Plant Science 172:433– 440.

9. Boujnane, M., Boukerb, A.M., Connil, N. Bacterial gene expression in response to catecholamine stress hormones. Curr Opin Endocr Metab Res 36:100543.

10. Boukerb AM, Cambronel M, Rodrigues S, Mesguida O, Knowlton R, Feuilloley MGJ, Zommiti M, Connil N. 2021. Inter-Kingdom Signaling of Stress Hormones: Sensing, Transport and Modulation of Bacterial Physiology. Front Microbiol 12:690942.

11. Martín-Mora D, Ortega Á, Matilla MA, Martínez-Rodríguez S, Gavira JA, Krell T. 2019. The Molecular Mechanism of Nitrate Chemotaxis via Direct Ligand Binding to the PilJ Domain of McpN. mBio 10:e02334–18.

12. Martín-Mora D, Ortega Á, Pérez-Maldonado FJ, Krell T, Matilla MA. 2018. The activity of the C4-dicarboxylic acid chemoreceptor of Pseudomonas aeruginosa is controlled by chemoattractants and antagonists. Sci Rep 8:2102.

13. Martín-Mora D, Ortega A, Reyes-Darias JA, García V, López-Farfán D, Matilla MA, Krell T. 2016. Identification of a Chemoreceptor in Pseudomonas aeruginosa That Specifically Mediates Chemotaxis Toward α-Ketoglutarate. Front Microbiol 7:1937.

14. Pasupuleti S, Sule N, Cohn WB, MacKenzie DS, Jayaraman A, Manson MD. 2014. Chemotaxis of Escherichia coli to norepinephrine (NE) requires conversion of NE to 3,4-dihydroxymandelic acid. J Bacteriol 196:3992–4000.

15. Bansal T, Englert D, Lee J, Hegde M, Wood TK, Jayaraman A. 2007. Differential effects of epinephrine, norepinephrine, and indole on Escherichia coli O157:H7 chemotaxis, colonization, and gene expression. Infect Immun 75:4597–607.

16. Gavira JA, Gumerov VM, Rico-Jiménez M, Petukh M, Upadhyay AA, Ortega A, Matilla MA, Zhulin IB, Krell T. 2020. How Bacterial Chemoreceptors Evolve Novel Ligand Specificities. mBio 11:e03066–19.

17. Corral-Lugo A, Matilla MA, Martín-Mora D, Silva Jiménez H, Mesa Torres N, Kato J, Hida A, Oku S, Conejero-Muriel M, Gavira JA, Krell T. 2018. High-Affinity Chemotaxis to Histamine Mediated by the TlpQ Chemoreceptor of the Human Pathogen Pseudomonas aeruginosa. mBio 9:e01894–18.

18. Xiang S-L, Xu K-Z, Yin L-J, Rao Y, Wang B, Jia A-Q. 2024. Dopamine, an exogenous quorum sensing signaling molecule or a modulating factor in Pseudomonas aeruginosa? Biofilm 8:100208.

19. Knecht LD, O’Connor G, Mittal R, Liu XZ, Daftarian P, Deo SK, Daunert S. 2016. Serotonin Activates Bacterial Quorum Sensing and Enhances the Virulence of Pseudomonas aeruginosa in the Host. EBioMedicine 9:161–169.

20. Medina Lopez AI, Fregoso DR, Gallegos A, Yoon DJ, Fuentes JJ, Crawford R, Kaba H, Yang H-Y, Isseroff RR. 2022. Beta adrenergic receptor antagonist can modify Pseudomonas aeruginosa biofilm formation in vitro: Implications for chronic wounds. FASEB J 36:e22057.

21. Cambronel M, Tortuel D, Biaggini K, Maillot O, Taupin L, Réhel K, Rincé I, Muller C, Hardouin J, Feuilloley M, Rodrigues S, Connil N. 2019. Epinephrine affects motility, and increases adhesion, biofilm and virulence of Pseudomonas aeruginosa H103. Sci Rep 9:20203.

22. Perraud Q, Kuhn L, Fritsch S, Graulier G, Gasser V, Normant V, Hammann P, Schalk IJ. 2022. Opportunistic use of catecholamine neurotransmitters as siderophores to access iron by Pseudomonas aeruginosa. Environ Microbiol 24:878–893.

23. Freestone PP, Lyte M, Neal CP, Maggs AF, Haigh RD, Williams PH. 2000. The mammalian neuroendocrine hormone norepinephrine supplies iron for bacterial growth in the presence of transferrin or lactoferrin. J Bacteriol 182:6091–6098.

24. Li J, Ma X, Zhao L, Li Y, Zhou Q, Du X. 2020. Extended Contact Lens Wear Promotes Corneal Norepinephrine Secretion and Pseudomonas aeruginosa Infection in Mice. Invest Ophthalmol Vis Sci 61:17.

25. Ma X, Wang Q, Song F, Li Y, Li J, Dou S, Xie L, Zhou Q. 2020. Corneal epithelial injury-induced norepinephrine promotes Pseudomonas aeruginosa keratitis. Exp Eye Res 195:108048.

