## Supplementary Figures and Methods for "*Pseudomonas aeruginosa* performs chemotaxis to serotonin, dopamine, epinephrine, and norepinephrine"

to

by

Elizabet Monteagudo-Cascales, Andrea Lozano-Montoya and Tino Krell

**Fig. S1) Chemotaxis response of *P. aeruginosa* and the Δ*pctABC*Δ*tlpQ* quadruple mutant towards 5 mM dopamine, epinephrine, serotonin, and norepinephrine.** Data are means and standard deviations from at least three biological replicates conducted in triplicate. Data have been corrected with the number of cells that swam into buffer-containing capillaries (4071 ± 1536 for wt and for 3127 ± 686 Δ*pctABCΔtlpQ* mutant).

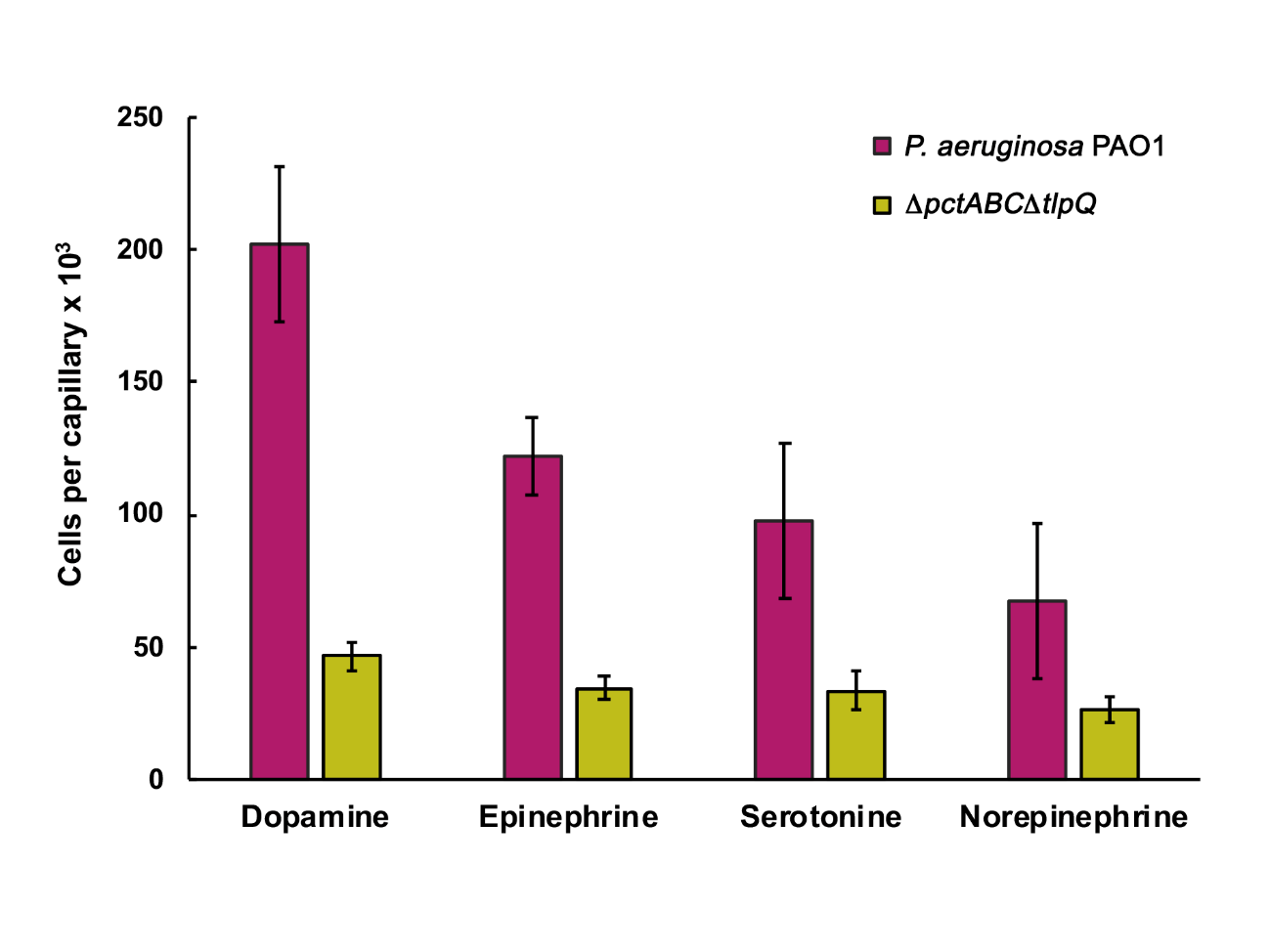

**Fig. S2) Quantitative capillary chemotaxis assay of *P. aeruginosa*, a mutant deficient in the *tlpQ* gene and Δ*tlpQ* mutant complemented with the p*tlpQ* plasmid towards 0.1 % (w/v) casamino acids.** Data are means and standard deviations from at least three biological replicates conducted in triplicate. Data have been corrected with the number of cells that swam into buffer-containing capillaries (2816 ± 724 for wt; 4440 ± 461 for Δ*tlpQ* and for 2109 ± 419 Δ*tlpQ*-p*tlpQ*).

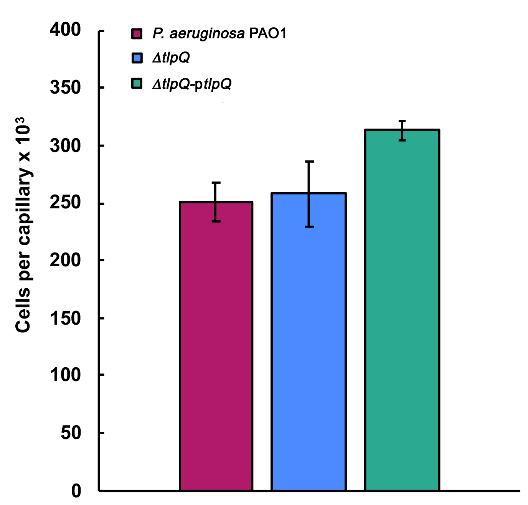

**Materials and Methods**

**Quantitative chemotaxis capillary assays**: Assays were conducted as reported in (4) with the exception that neurotransmitter solutions were prepared immediately before use and protected against light. Assays were conducted under dim light conditions. The construction of the *tlpQ* deletion mutant (PAO1 Δ*tlpQ*) and the Δ*pctABCΔtlpQ* quadruple mutant (PCT2Q) has been reported in (2). The construction of the plasmid pTlpQ for mutant complementation has been reported in (3). The plasmid was introduced into PAO1 Δ*tlpQ* by electroporation. The resulting strain was selected by growth on LB plates containing 500 μg ml^−1^ carbenicillin.

**Protein overexpression and purification**: TlpQ-LBD was overexpressed and purified as reported in (2).

**Isothermal titration calorimetry**: Experiments were conducted on a VP-microcalorimeter (Microcal, Amherst, MA, USA) at 25 ºC (for dopamine) and at 10 ºC (for remaining compounds). Freshly purified protein was dialyzed into 3 mM Tris, 3 mM PIPES, 3 mM MES, 150 mM NaCl, 10 % (v/v) glycerol, pH 7.0, adjusted to a concentration of 18 to 80 μM and titrated with 0.2 to 10 mM ligand solutions made up in dialysis buffer. Typically, a single injection of 1.6 µl was followed by a series of 4.8 to 14.4 µl aliquots. For competition assays, serotonin and norepinephrine were added to the protein (sample cell) and the spermidine solution (injector syringe) at a final concentration of 10 mM. The mean enthalpies measured from the injection of ligand solutions into the buffer were subtracted from raw titration data. Data were normalized with the ligand concentrations, the first data point removed and the data fitted with the ‘One Binding Site’ model of the MicroCal version of ORIGIN (Microcal, Amherst, MA, USA).

**References**

1.	Martín-Mora D, Ortega A, Reyes-Darias JA, García V, López-Farfán D, Matilla MA, Krell T. 2016. Identification of a Chemoreceptor in *Pseudomonas aeruginosa* That Specifically Mediates Chemotaxis Toward α-Ketoglutarate. Front Microbiol 7:1937.

2.	Corral-Lugo A, Matilla MA, Martín-Mora D, Silva Jiménez H, Mesa Torres N, Kato J, Hida A, Oku S, Conejero-Muriel M, Gavira JA, Krell T. 2018. High-Affinity Chemotaxis to Histamine Mediated by the TlpQ Chemoreceptor of the Human Pathogen *Pseudomonas aeruginosa*. mBio 9:e01894-18.

3.	Kim HE, Shitashiro M, Kuroda A, Takiguchi N, Kato J. 2007. Ethylene chemotaxis in *Pseudomonas aeruginosa* and other *Pseudomonas* species. Microbes Environ 22:186–189.

4.	Matilla MA, Velando F, Tajuelo A, Martin-Mora D, Xu W, Sourjik V, Gavira JA, Krell T. 2022. Chemotaxis of the human pathogen *Pseudomonas aeruginosa* to the neurotransmitter acetylcholine. mBio e0345821.
